# Directed evolution for improved total secretory protein production in *Escherichia coli*

**DOI:** 10.1101/2020.11.03.366773

**Authors:** David Gonzalez-Perez, James Ratcliffe, Shu Khan Tan, Mary Chen May Wong, Yi Pei Yee, Natsai Nyabadza, Jian-He Xu, Tuck Seng Wong, Kang Lan Tee

**Affiliations:** Department of Chemical & Biological Engineering, The University of Sheffield, Sir Robert Hadfield Building, Mappin Street, Sheffield S1 3JD, UK; Moffitt Cancer Center & Research Institute, Department of Drug Discovery, Stabile Research Building, 12902 Magnolia Dr, Tampa, FL 33612, USA; Laboratory of Biocatalysis and Bioprocessing, State Key Laboratory of Bioreactor Engineering, East China University of Science and Technology, 130 Meilong Road, Shanghai 200237, People’s Republic of China; National Center for Genetic Engineering and Biotechnology, 113 Thailand Science Park, Phahonyothin Road, Khlong Luang, Pathum Thani 12120, Thailand

**Author notes:** Address correspondence to: **Dr. Kang Lan Tee**, Or, **Dr. Tuck Seng Wong**.

**Keywords:** Secretory protein production, protein secretion, directed evolution, protein engineering, OsmY, carrier protein, *Escherichia coli*

## Abstract

Production of secretory protein in Gram-negative bacteria simplifies downstream processing in recombinant protein production, accelerates protein engineering, and advances synthetic biology. Signal peptides and secretory carrier proteins are commonly used to effect the secretion of heterologous recombinant protein in Gram-negative bacteria. The *Escherichia coli* osmotically-inducible protein Y (OsmY) is a carrier protein that secretes a target protein extracellularly, and we have successfully applied it in the <u>B</u>acterial <u>E</u>xtracellular Protei<u>n</u> Secretio<u>n</u> S<u>y</u>stem (BENNY) to accelerate the directed evolution workflow. In this study, we applied directed evolution on OsmY to enhance its total secretory protein production.

After just one round of directed evolution followed by combining the mutations found, OsmY(M3) (L6P, V43A, S154R, V191E) was identified as the best carrier protein. OsmY(M3) produced 3.1 ± 0.3 fold and 2.9 ± 0.8 fold more secretory Tfu0937 β-glucosidase than its wildtype counterpart in *E. coli* strains BL21(DE3) and C41(DE3), respectively. OsmY(M3) also produced more secretory Tfu0937 at different cultivation temperatures (37 °C, 30 °C and 25 °C). Subcellular fractionation of the expressed protein confirmed the essential role of OsmY in protein secretion. Up to 80.8 ± 12.2% of total soluble protein was secreted after 15 h of cultivation. When fused to a red fluorescent protein or a lipase from *Bacillus subtillis*, OsmY(M3) also produced more secretory protein compared to the wildtype.

This is the first report of applying directed evolution on a carrier protein to enhance total secretory protein production. The methodology can be further extended to evolve other signal peptides or carrier proteins for secretory protein production in *E. coli* and other bacteria. In this study, OsmY(M3) improved the production of three proteins, originating from diverse organisms and with diverse properties, in secreted form, clearly demonstrating its wide-ranging applications.

## INTRODUCTION

### Protein secretion in Gram-negative bacteria

Recombinant protein production, including technical enzymes and biopharmaceutical proteins, is a growing multi-billion-dollar market. Different eukaryotic and prokaryotic hosts are currently used for recombinant protein production, including mammalian cells, fungi, yeast and bacteria. Bacteria are especially attractive for their fast growth rates, ease in handling, broad range of carbon sources and relatively low-cost culture medium. Well-established and widely accessible genetic manipulation tools are also available, especially for the Gram-negative bacterium *Escherichia coli* ^[1]^.

Gram-positive bacteria can secrete native proteins or proteins that originate from close relatives at high yield ^[2]^. However, their compatibility with other proteins is low and the presence of extracellular proteolytic degradation is a challenge for product quality assurance. In contrast, Gram-negative bacteria produce a broad range of heterologous proteins. Protein secretion in Gram-negative bacteria is less common compared to other organisms like Gram-positive bacteria, yeast and fungi. However, it offers several advantages. First, Gram-negative bacteria, like *E. coli*, secretes few native proteins to provide a relatively pure background for heterologous protein production. Second, no cell disruption is required, hence reducing the complexity and the cost of downstream processing. Third, protein secretion out of the cytoplasm provides an oxidizing environment that facilitates disulphide bond formation and protein folding, for example during hormones and antibody fragments production. Fourth, toxic effect of some target proteins on the production host can be alleviated when the protein is secreted out of the cell.

Protein secretion in Gram-negative bacteria can be achieved by fusing a signal peptide or a secretory carrier to the N- or C-terminus of the target protein ^[3, 4]^. Enhancing protein secretion is a challenge as it is currently not possible to predict the performance of a signal peptide or a secretory carrier for a specific target protein or host. Three approaches are generally used for secretory protein production in *E. coli*. The first approach is to screen a large library of signal peptides and secretory carriers ^[5, 6]^ to identify a fusion partner for optimal secretion. The second approach is to engineer the signal peptides or the transporter proteins for a better extracellular heterologous protein production in *E. coli*. Examples include: (i) inserting short random peptides libraries (1 to 13 amino acids) at the junction between the signal peptide and the mature protein followed by screening for the construct that secreted most protein to the *E. coli* periplasm ^[7]^, (ii) screening error-prone mutagenesis library of an ABC transporter from *Pseudomonas fluorescens* to enhance its extracellular secretion efficiency in *E. coli* ^[8]^, (iii) directed evolution of translation initiation region without altering the amino acid sequence of signal peptide to enhance protein secretion into the *E. coli* periplasm ^[9]^, and (iv) using non-optimal codons in the signal sequence of TolB for a better extracellular protein secretion in *E. coli* ^[10]^. The third approach involves engineering the protein production host, for instance, perturbing the cell wall peptidoglycan network via deletion of *dacA* and *dacB* ^[11]^ and screening *E. coli* single-gene knockout library for hypersecretory phenotypes ^[12]^.

### The osmotically-inducible protein Y from *Escherichia coli*

The *E. coli* osmotically-inducible protein Y (OsmY) is expressed in response to a variety of stress conditions, especially osmotic shock ^[13–15]^. It was identified as a naturally excreted protein in a systematic proteomic analysis of the extracellular proteome of *E. coli* BL21(DE3) ^[16]^. OsmY has two BON (bacterial OsmY and nodulation) domains; one between amino acids 55-123 and the second between amino acids 134-201. The BON domain is a conserved domain that has typically about 60 amino acid residues, and has an α/β predicted fold. The exact function of the BON domain is unclear, but it is predicted to be a phospholipid binding domain that explains OsmY’s secretory function ^[24]^. Numerous studies used OsmY for extracellular production of recombinant proteins in *E. coli*, including endoglucanase ^[17]^, β-glucosidase ^[17]^, xylanase ^[18, 19]^, xylosidase ^[18, 19]^, single-chain antibody ^[20]^ and various human proteins ^[21]^. We also incorporated OsmY in the <u>B</u>acterial <u>E</u>xtracellular Protei<u>n</u> Secretio<u>n</u> S<u>y</u>stem (BENNY) to accelerate the directed evolution workflow in engineering DyP4, a heme peroxidase ^[22]^. OsmY was previously reported to show chaperone activity and helped optimized unstable protein folding *in vivo* ^[23]^. The versatility of OsmY prompted us to engineer this protein to further expand its application.

In this study, we demonstrated the first successful example of directed evolution on a secretory carrier to enhance total secretory protein production in *E. coli*. There are two mechanisms by which total secretory protein production can be enhanced: (a) increasing the secretion capability of a carrier (*i.e.*, altering protein partition by favouring extracellular secretion), and (b) increasing the expression level of the target protein carrying a secretion signal such as a carrier. OsmY is strategically chosen, as it has the potential to satisfy either or both requirements.

## RESULTS AND DISCUSSION

### Directed evolution of OsmY

To evolve OsmY for a higher total secretory protein production, β-glucosidase Tfu0937 from *Thermobifida fusca* YX was selected as the secretion target protein for its validated application in cellulose degradation and ease of monitoring enzymatic acitivty. OsmY, the secretory carrier, was fused to the N-terminus of Tfu0937 and cloned into a pET24a(+) vector to create the plasmid pET24a-OsmY-Tfu0937 (Fig. S1). Activity of secreted OsmY-Tfu0937 was monitored using the *p*-nitrophenyl-β-D-glucopyranoside (pNPG) assay, where the absorbance of the *p*-nitrophenolate released was measured at 405 nm. OsmY can secrete proteins in different *Escherichia coli* strains ^[16]^. To select the more efficient strain for subsequent directed evolution, the levels of total secreted OsmY-Tfu0937 produced by *E. coli* C41(DE3) and BL21(DE3) were first measured using the pNPG assay. C41(DE3) cultivation gave higher OsmY-Tfu0937 activity in the spent medium compared to BL21(DE3), and was therefore chosen for subsequent directed evolution experiments. The pNPG assay was adapted to a 96-well format. The final optimised assay had an apparent coefficient of variance of 19.7% (Fig. 1a), sufficient for a high throughput screening. The enzymatic activity of secreted OsmY-Tfu0901 using pNPG as a pseudosubstrate was also directly proportional to the volume of spent medium, when 5–50 μL was used (Fig. 1b). Variable volume of spent medium was thus used to keep the assay within the linear detection range, especially for OsmY variants with higher total secretory protein production.

**Figure 1:**
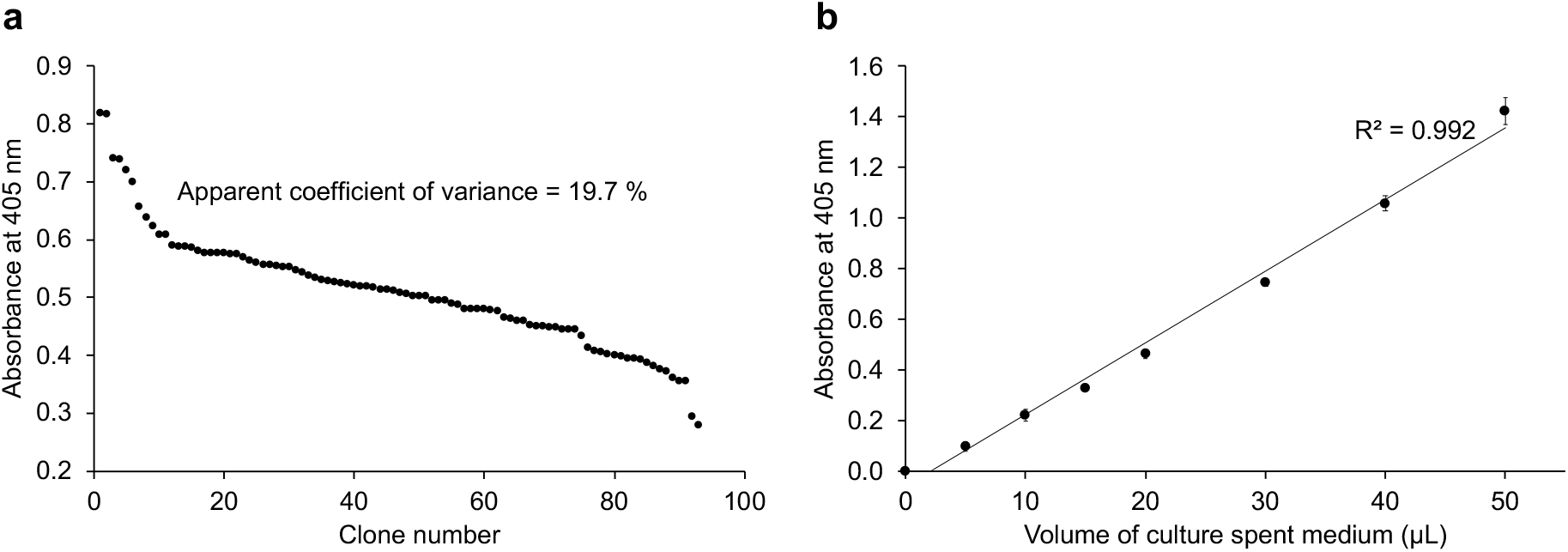
The OsmY(WT)-Tfu0937 activity in the spent medium was measured with the pNPG assay using an absorbance at 405 nm. (**a**) The pNPG assay was performed in a 96-well microplate for OsmY(WT)-Tfu0937-expressing C41(DE3) cells cultivated in a 96-well deep well plates. The coefficient of variance was calculated using the absolute values measured at 405 nm. (**b**) The pNPG assay was conducted using 5 – 50 μL of spent medium. The activity value is linear to the volume of spent medium used with an R^2^ = 0.992.

Error-prone PCR (epPCR) was used to prepare the random mutagenesis libraries. Mutations were only targeted to the sequence encoding for OsmY, leaving the Tfu0937 sequence unaltered. This was designed to ensure all improvements can be attributed to OsmY and were not due to changes in the target protein, Tfu0937. The libraries were created using two different conditions with high and low mutagenic rates (Fig. 2). The error rate of Taq DNA polymerase was manipulated by increasing Mg^2+^ concentration, adding Mn^2+^, applying imbalanced nucleotide concentration or utilizing combinations of these factors ^[22, 24]^. The effects of different mutagenic rates were evident in the activity of the clones in the library. For the low mutagenic rate library (Fig. 2a), 31% of clones had the same activity as the parent (*i.e.*, one standard deviation above and below the average of parental clones) and 30% of clones were more active than the parent (*i.e.*, greater than one standard deviation above the parent average). For the high mutagenic rate library (Fig. 2b), 16% of clones had the same activity as the parent and 21% of clones were more active than the parent. More inactive clones were also found in the high mutagenic rate library (23%) compared to the low mutagenic rate library (5%).

**Figure 2:**
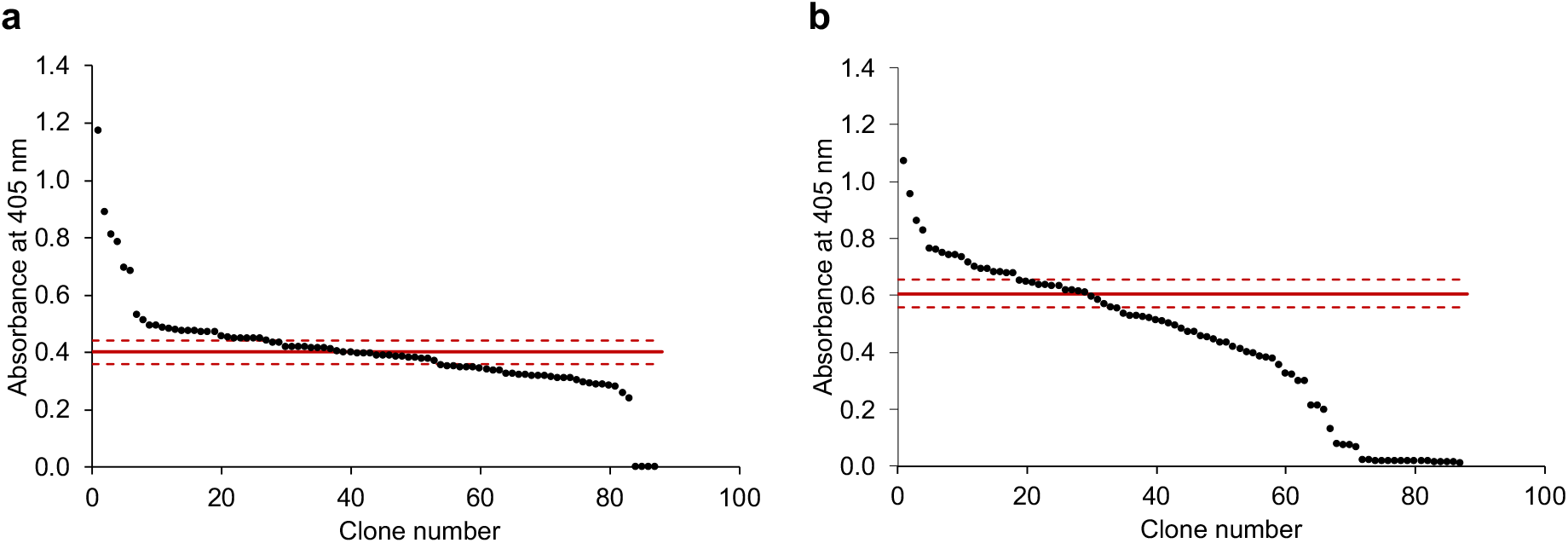
The OsmY-Tfu0937 activity in the spent medium of epPCR generated OsmY mutagenesis libraries was measured in a 96-well microplate. The activity was measured with the pNPG assay using an absorbance at 405 nm. The libraries were generated using (**a**) a low and (**b**) a high mutagenic rates. The activities of parental clones in column 6 were measured in the same 96-well microplate. The average of parental clones is shown as a solid red line and one standard deviation above and below the average are shown as dash red lines.

### Improved OsmY mutants obtained from both random mutagenesis libraries

About three hundred clones from the two random mutagenesis libraries were screened and four improved variants were identified (Table 1). These four variants were subsequently cultivated in 5-mL tube cultures and extracellular OsmY-Tfu0937 activity was measured again to confirm their improvements as shown in Table 1. Mutations in these variants were verified by DNA sequencing. The variants and wildtype can be arranged in decreasing order of extracellular OsmY-Tfu0937 production as OsmY(TOA4) > OsmY(S19C) > OsmY(A39) > OsmY(V43A) > OsmY(WT). In agreement with the mutagenic conditions, the three variants from the low mutagenic rate library have a single nucleotide substitution each, while the one variant from the high mutagenic rate library has four nucleotide substitutions.

**Table 1:** OsmY variants identified from screening of epPCR libraries. The mutations were verified by DNA sequencing.

| Mutagenic rate | Variant name | Missense mutation | Silent mutation | Relative activity compared to OsmY wildtype* |
| --- | --- | --- | --- | --- |
| n.a. | OsmY(WT) | n.a. | n.a. | 1.00 ± 0.08 |
| Low | OsmY(S19C) | S19C (A → T) | n.a. | 3.40 ± 0.25 |
| Low | OsmY(V43A) | V43A (T → C) | n.a. | 1.41 ± 0.13 |
| Low | OsmY(A39) | n.a. | A39 (G → T) | 1.67 ± 0.16 |
| High | OsmY(TOA4) | L6P (T → C) | K107 (A → G) | 5.50 ± 0.56 |
|  |  | S154R (A → C) |  |  |
|  |  | V191E (T → A) |  |  |

\*Variants were cultivated in 5-mL tubes cultures and activity of extracellular OsmY-Tfu0937 was determined with the pNPG assay using an absorbance at 405 nm, and compared with the wildtype OsmY-Tfu0937.

OsmY is a 201-amino acid protein with an N-terminal 28-amino acid signal peptide. Interestingly, the top two variants have mutations in their signal peptides; L6P for OsmY(TOA4) and S19C for OsmY(S19C). It was previously shown that the OsmY signal peptide secreted protein into the periplasm, but the full length OsmY protein was required for extracellular protein secretion ^[16]^. Signal peptides have a conserved tripartite overall structure, which consists of a positively charged amino-terminal region, a central hydrophobic core, and a polar carboxyl-terminal region ^[25]^. The mutations L6P and S19C conserved the respective hydrophobic and polar nature of the amino acids. Variant OsmY(A39) has a silent mutation that changes the codon for A39 from GCG→GCT. Interestingly, codon GCG has the highest usage frequency among the four codons for alanine while GCT has the lowest usage frequency. The same was observed for the K107 silent mutation in OsmY(TOA4), where the higher usage frequency codon (AAA) was mutated to the lower usage frequency codon (AAG) for lysine. Non-optimal codon in the signal peptide had been reported to slow down the kinetics of translation to positively impact secretion efficiency ^[10]^. The enhanced production of secretory protein by a switch to less optimal codons in A39 and K107 could potentially also be due to a slower translation kinetics. Important to note, codon deoptimization of signal peptides also improved the production of secretory protein in baculovirus/insect cell expression system ^[26]^. Our directed evolution data, along with the results from other signal peptide engineering studies, allow us to propose engineering strategies to guide future improvement of secretory protein production across various protein expression systems: (a) preserving the tripartite nature of the signal peptides through conservative mutations, and (b) slowing the protein translation by deoptimizing the codons of the carrier protein.

### Variant from directed evolution produced 1.8-times more extracellular protein

The three variants with missense mutations [OsmY(TOA4), OsmY(S19C), and OsmY(V43A)] were re-transformed into C41(DE3) to rule out any possible effects due to host strain instability, before they were cultivated in 50-mL flask cultures at 37 °C for further characterization. Secreted protein was detected 5.5 h after culture inoculation, however significant activity was observed after 24 h of cultivation. Results (Fig. 3a) showed that variant OsmY(TOA4) had 1.8 ± 0.5 times more OsmY-Tfu0937 activity in the spent medium compared to the OsmY(WT) after 24 h cultivation, and this improvement increased further to 2.2 ± 0.3 times after 48 h. Enhanced secretory protein production did not affect the cell viability, reflected by the similar cell growth of *E. coli* expressing OsmY(WT)-Tfu0937 and the variants.

**Figure 3:**
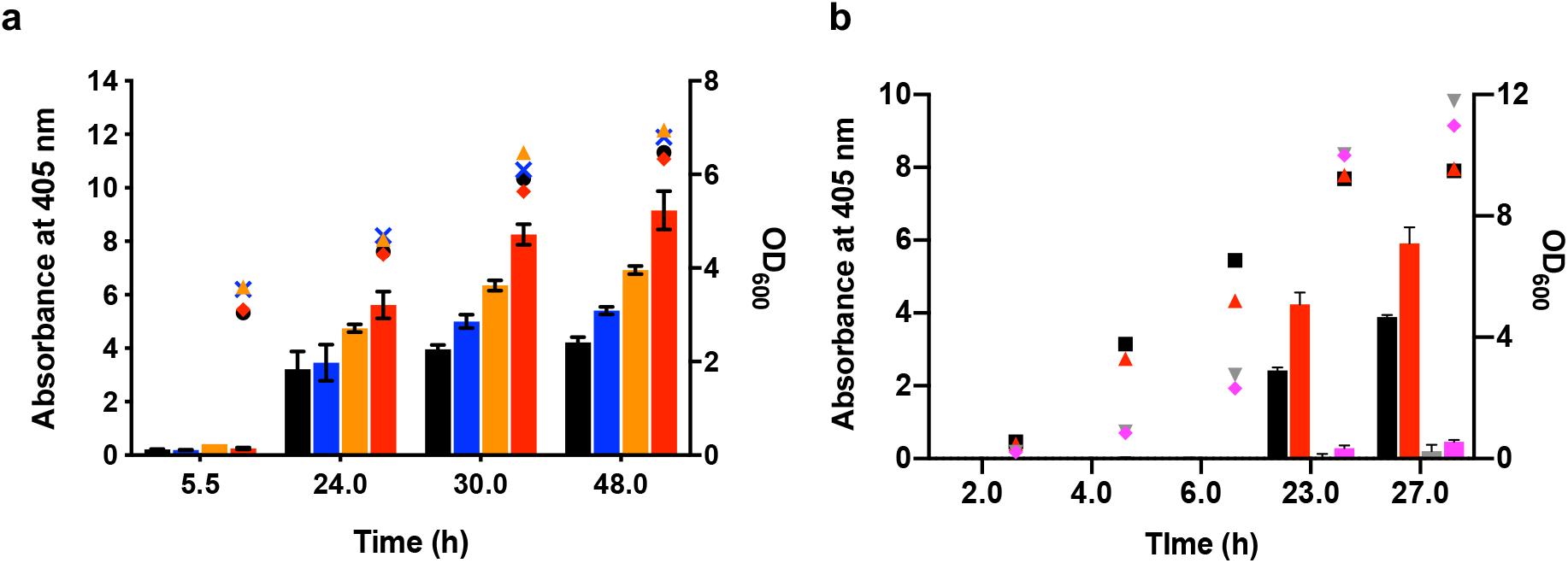
Characterization of improved OsmY variants in 50-mL flask cultures. The OsmY-Tfu0937 activity, measured with the pNPG assay using an absorbance at 405 nm (left axis), are plotted as bars. The cell optical densities monitored at 600 nm (right axis) are shown as symbols. Cultures were inoculated using 1:100 dilution. (**a**) OsmY(WT) and variants were expressed using C41(DE3) in 2×TY auto-induction media and at 37 °C and 200 rpm. The bars and symbols of the variants are represented using different colours; OsmY(WT) (black), OsmY(V43A) (blue), OsmY(S19C) (orange) and OsmY(TOA4) (red). Experiments were performed in triplicate. (**b**) OsmY(WT)-Tfu0937 and OsmY(TOA4)-Tfu0937 were expressed using C41(DE3) in 2×TY auto-induction media, at 30 °C or 25 °C, and 200 rpm. The bars and symbols of the sample are represented using different colours; OsmY(WT) at 30 °C (black), OsmY(TOA4) at 30 °C (red), OsmY(WT) at 25 °C (grey), OsmY(TOA4) at 25 °C (magenta).

Tfu0937 secretion by OsmY(WT) and OsmY(TOA4) were also compared at cultivation temperatures of 25 °C and 30 °C (Fig. 3b). OsmY(TOA4) was more efficient than wildtype at these lower temperatures in secretory protein production. When cultivated at 25 °C, OsmY-Tfu0937 activity in the spent medium remained low throughout the cultivation, likely due to the slower growth and protein production rates compared to cultivation at higher temperatures. However, more extracellular activity was detected for the OsmY(TOA4) variant than the OsmY(WT) after 27 h. When cultivated at 30 °C, OsmY-Tfu0937 activity in the spent medium of OsmY(TOA4) was 1.8 ± 0.2 times (23.0 h) and 1.5 ± 0.1 times (27.0 h) that of the wildtype, similar to the improvement seen at 37 °C cultivation (Fig. 3a). As in the 37 °C cultivation, there was no significant difference in cell growth between OsmY(TOA4) and OsmY(WT) at 30 °C or at 25 °C. However, higher final cell optical densities were obtained at lower temperatures, potentially due to lower protein production load.

### Combination of mutations further enhanced total secretory protein production

Site directed mutagenesis was used to add the mutations S19C and V43A individually to the best variant OsmY(TOA4), creating mutants OsmY(M1) (*i.e.*, TOA4+S19C) and OsmY(M3) (*i.e.*, TOA4+V43A), respectively. When cultivated at the same conditions used in Fig. 3a, the pNPG assay showed that OsmY(M3) resulted in 2.9 ± 0.8 times more secretory protein compared to OsmY(WT) at 24 h (Fig. S2). As with the earlier experiments, cell growth of variant OsmY(M3) was similar to that of OsmY(WT). Contrary to OsmY(M3), the spent medium of variant OsmY(M1) showed no OsmY-Tfu0937 activity, though its cell growth was similar to OsmY(WT). To determine if OsmY(M1) had loss only its protein secretion capability or also protein expression, enzymatic lysis with lysozyme was used to extract the intracellular soluble protein in OsmY(M1). The extracted soluble protein showed no OsmY-Tfu0937 activity when measured using the pNPG assay, suggesting the loss of soluble OsmY-Tfu0937 expression in variant OsmY(M1). We subsequently also added S19C to variant OsmY(M3) and observed the same effect, where no secreted OsmY-Tfu0937 was detected by the pNPG assay. We therefore conclude that combining S19C with the existing mutations in OsmY(TOA4) led to negative epistatic interaction and loss of soluble protein expression.

### L6P is the beneficial mutation in OsmY(TOA4) variant

OsmY(TOA4) was the best variant identified from our initial screening. It has three missense mutations (L6P, S154R, V191E), with one located in the signal peptide (L6P) and the remaining two in the second BON domain. A single mutant was created for each of these mutations using site-directed mutagenesis and their extracellular OsmY-Tfu0937 activity was measured to deconvolute their effects on secretory protein production (Fig. 4). The two single mutants, S154R and V191E, showed the same OsmY-Tfu0937 activity in the spent medium as the wildtype OsmY. However, the L6P single mutant showed 3.1 ± 1.6 times the level of secreted protein compared to the wildtype OsmY after 24.0 h cultivation. Comparing the secretory protein production levels achieved by OsmY(TOA4) and OsmY(L6P), 1.8 ± 0.5 times and 3.1 ± 1.6 times respectively, both variants secreted similar amount of proteins. There was no additive interaction between S154R or V191E with L6P to further increase secretory protein production in OsmY(TOA4). The cell growth of the three single mutants were similar to the OsmY(WT), as was observed for OsmY(TOA4) (Fig. S3). Our data indicated that the signal peptide of OsmY plays a greater role in secretory protein production compared to its BON domain.

**Figure 4:**
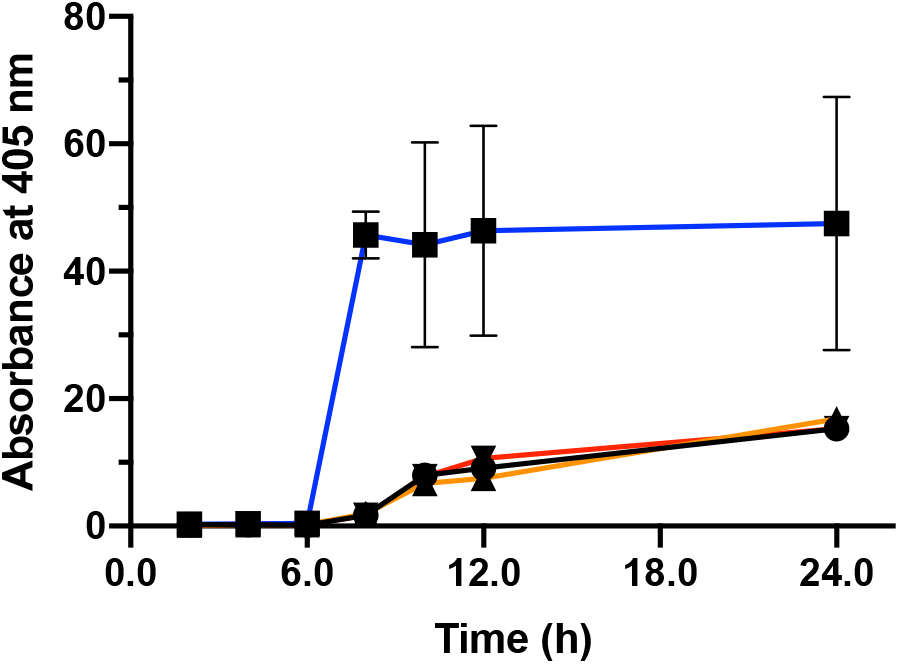
The OsmY-Tfu0937 activity in the spent medium of single site-directed mutants was measured with the pNPG assay using an absorbance at 405 nm. Cells were cultivated in 20-mL of 2×TY auto-induction media at 37 °C. Culture was inoculated using 1:100 dilution and 0.5 mL culture was sampled at various time points for activity assay. Experiments were performed in triplicate. The samples are represented using different colours; OsmY(WT) – black line, OsmY(L6P) – blue line, OsmY(S154R) – orange line and OsmY(V191E) – red line.

### OsmY is required for extracellular protein secretion in *E. coli*

Tfu0937 was found in the secretome of *T. fusca* ^[27]^. To confirm that OsmY is necessary for the secretion of Tfu0937 in *E. coli* and to rule out the possibilities that this is solely due to cell leakage or the presence of a secretion motif in Tfu0937 recognized by *E. coli*, the plasmid pET24a-Tfu0937 was constructed. Plasmids pET24a-Tfu0937 and pET24a-OsmY(M3)-Tfu0937 were used for protein expression. Following a 15-h cultivation, the extracellular, periplasmic and cytoplasmic fractions were extracted and Tfu0937 activities in individual fractions were quantified using the pNPG assay. When Tfu0937 was expressed without OsmY using plasmid pET24a-Tfu0937, the extracellular Tfu0937 activity constituted only 1.8 ± 0.3% of the total Tfu0937 activity in the three fractions (Fig. 5). Most of the Tfu0937 activity was found in the periplasmic fraction while the cytoplasmic fraction accounted for 18.8 ± 3.1% of total Tfu0937 activity. In contrast, 80.8 ± 12.2% of total OsmY-Tfu0937 activity was found in the spent medium when OsmY(M3) was fused to the N-terminus of Tfu0937 (Fig. 5).

**Figure 5:**
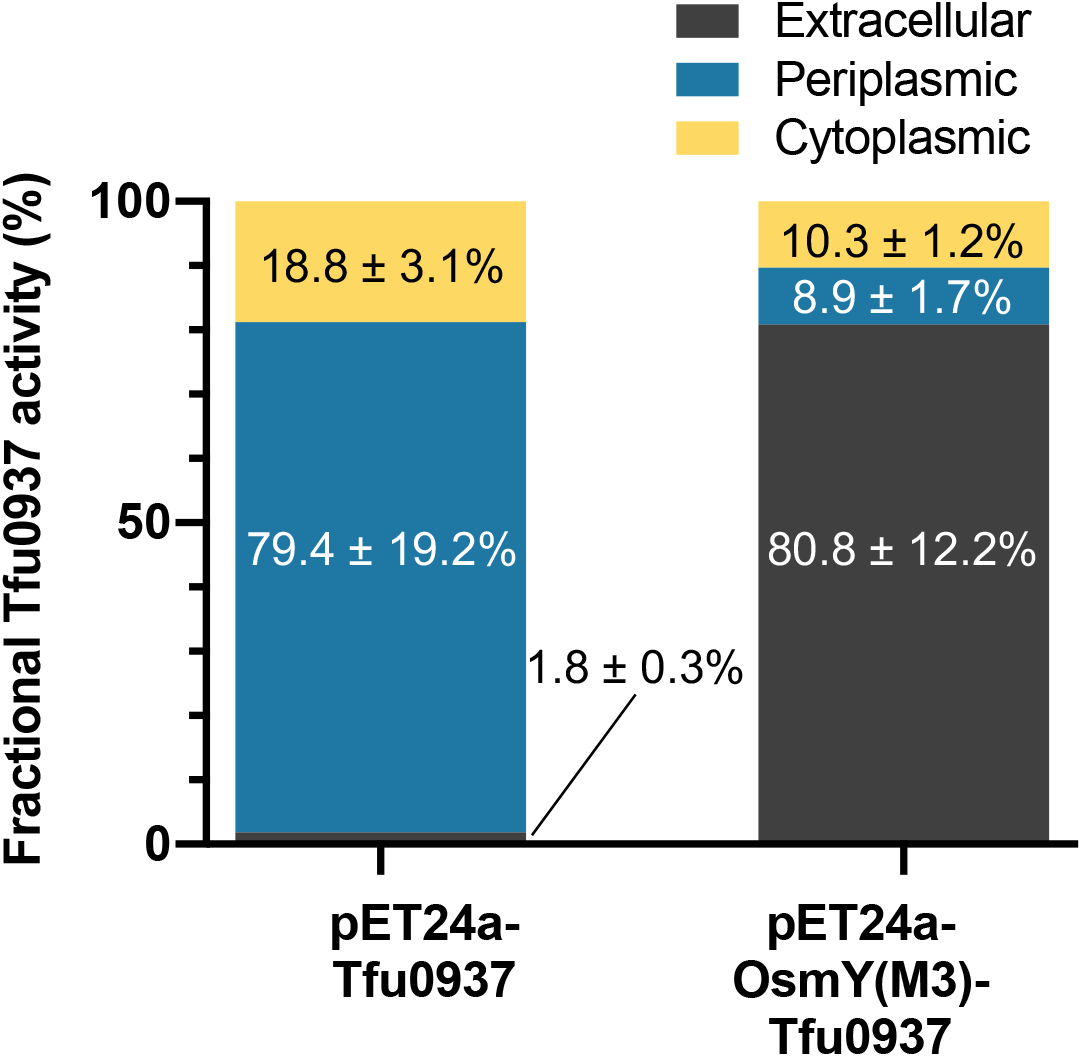
Subcellular localization of Tfu0937 and OsmY(M3)-Tfu0937. Proteins were expressed using pET24a-Tfu0937 and pET24a-OsmY(M3)-Tfu0937 in 2×TY auto-induction medium at 37 °C for 15 h. The fractional Tfu0937 and OsmY(M3)-Tfu0937 activity in the extracellular, periplasmic and cytoplasmic fractions were measured with the pNPG assay using an absorbance at 405 nm. The enzyme activity in each fraction was calculated as a percentage of total activity measured in all three fractions. Experiments were performed in triplicate.

### OsmY(M3) variant increased the total protein production

The increased extracellular OsmY-Tfu0937 observed in OsmY(M3) compared to wildtype can be attributed to one or both of these factors: (1) higher fractional secretion of total protein, and (2) increase in total protein expression. To understand the reason behind higher secretory protein production by OsmY(M3), we performed subcellular fractionation for OsmY(WT) and OsmY(M3) at two different time points during cultivation (6 h and 24 h). After 6 h cultivation, when cells were in the exponential growth phase, both wildtype and OsmY(M3) showed the most OsmY-Tfu0937 activity in the cytoplasmic fraction, followed by the periplasmic fraction and the least activity was detected in the secreted fraction (Fig. 6). The wildtype strain has a higher fraction of OsmY-Tfu0937 found extracellularly and in the periplasm compared to OsmY(M3). However, the OsmY(M3) variant had 3.64 ± 0.13 times more total OsmY-Tfu0937 activity compared to OsmY(WT). After 24 h cultivation, both wildtype and variant showed similar protein distribution in the extracellular, periplasmic and cytoplasmic fractions (Fig. 6). However, 2.73 ± 0.24 times more total OsmY-Tfu0937 activity was measured in OsmY(M3) compared to the wildtype. The enhanced production of secretory protein in OsmY(M3) was thus largely attributed to higher OsmY-Tfu0937 expression in the variant, while its secretion capability is retained.

**Figure 6:**
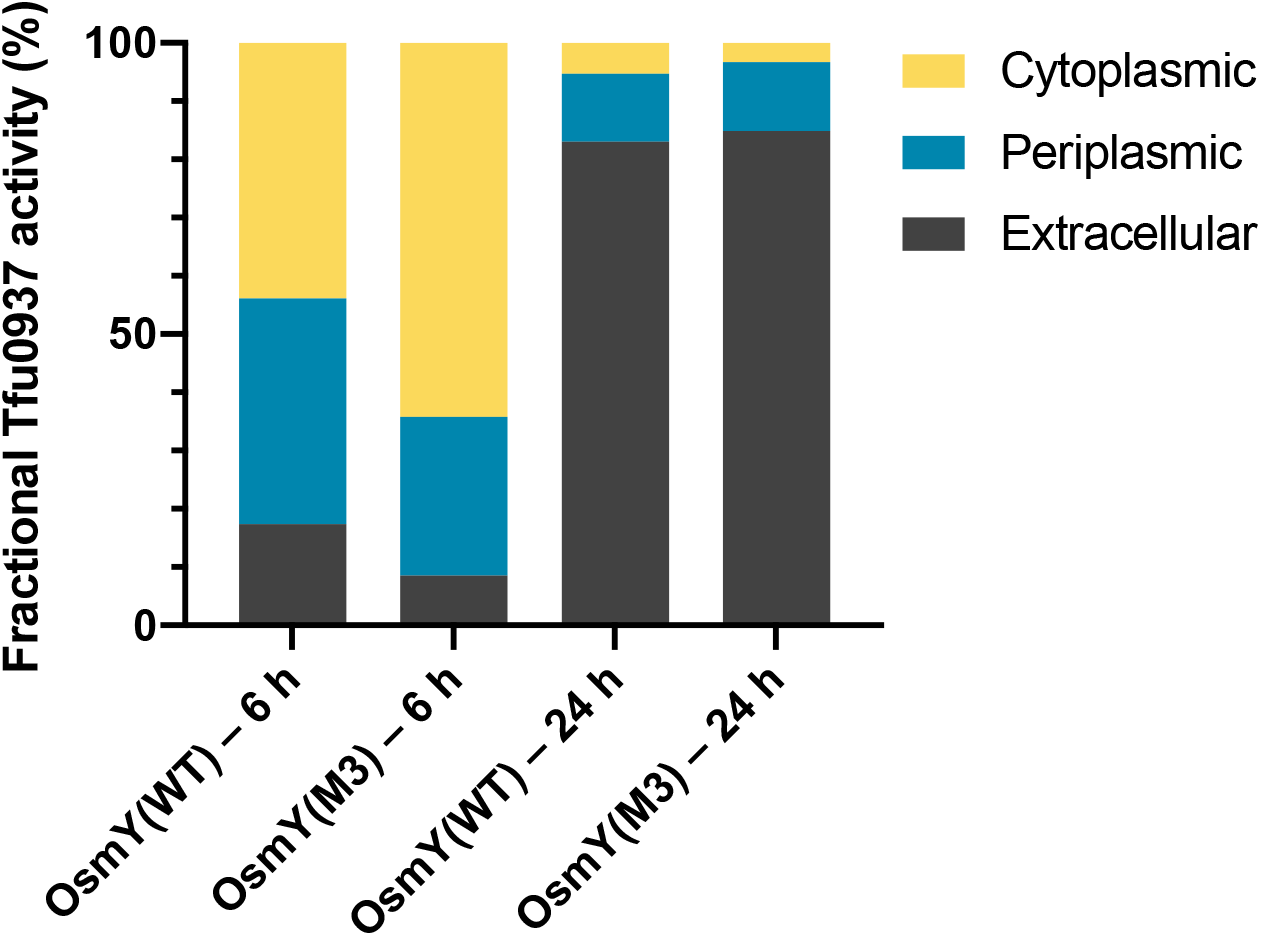
Fractional OsmY-Tfu0937 activity in the extracellular, periplasmic and cytoplasmic fractions of cells expressing Tfu0937 fused to either OsmY(WT) or OsmY(M3). Activity was measured with the pNPG assay using an absorbance at 405 nm. Experiments were performed in triplicate. Cells were cultivated in 2×TY auto-induction medium at 37 °C for either 6 h or 24 h. The enzyme activity in each fraction was calculated as a percentage of total activity in all three fractions.

### OsmY(M3) variant improves the production of secretory protein in both BL21(DE3) and C41(DE3) strains

The *E. coli* strain C41(DE3) was used for directed evolution, where OsmY(TOA4) was obtained and OsmY(M3) was subsequently derived. Since OsmY(WT) can secrete OsmY-Tfu0937 in both *E. coli* BL21(DE3) and C41(DE3) strains, we wanted to determine if the enhanced secretory protein production can be transferred to BL21(DE3). The OsmY-Tfu0937 secreted by OsmY(TOA4) and OsmY(M3) in BL21(DE3) were measured using the pNPG assay (Fig. 7). All variants showed similar cell growth over time in both C41(DE3) and BL21(DE3) strains. Protein secretion started in the late logarithmic phase of cell growth and most protein secretion occurred during the stationary phase. In agreement with initial experiments, the C41(DE3) strain produced more secreted protein compared to BL21(DE3), evident in the higher OsmY-Tfu0937 activity detected for the C41(DE3) host (Fig. 7a). Both OsmY(TOA4) and OsmY(M3) increased secretory protein level in BL21(DE3) compared to the wildtype. However, their improvements in BL21(DE3) were similar (Fig 7b). After 25 h of cultivation, OsmY(TOA4) and OsmY(M3) in BL21(DE3) cells produced 2.9 ± 0.3 times and 3.1 ± 0.3 times more secreted protein respectively when compared to the wildtype. This was different in C41(DE3) where OsmY(M3) produced more secreted protein than OsmY(TOA4) (Fig 7a). The improvements observed for OsmY(M3) were similar in both C41(DE3) and BL21(DE3) but OsmY(TOA4) showed greater improvement in BL21(DE3) compared to C41(DE3). We thus concluded that the improved secretory protein production by the OsmY variants in C41(DE3) can be transferred to BL21(DE3), though the mutations in OsmY(TOA4) likely affected secretion and/or expression in the two strains differently.

**Figure 7:**
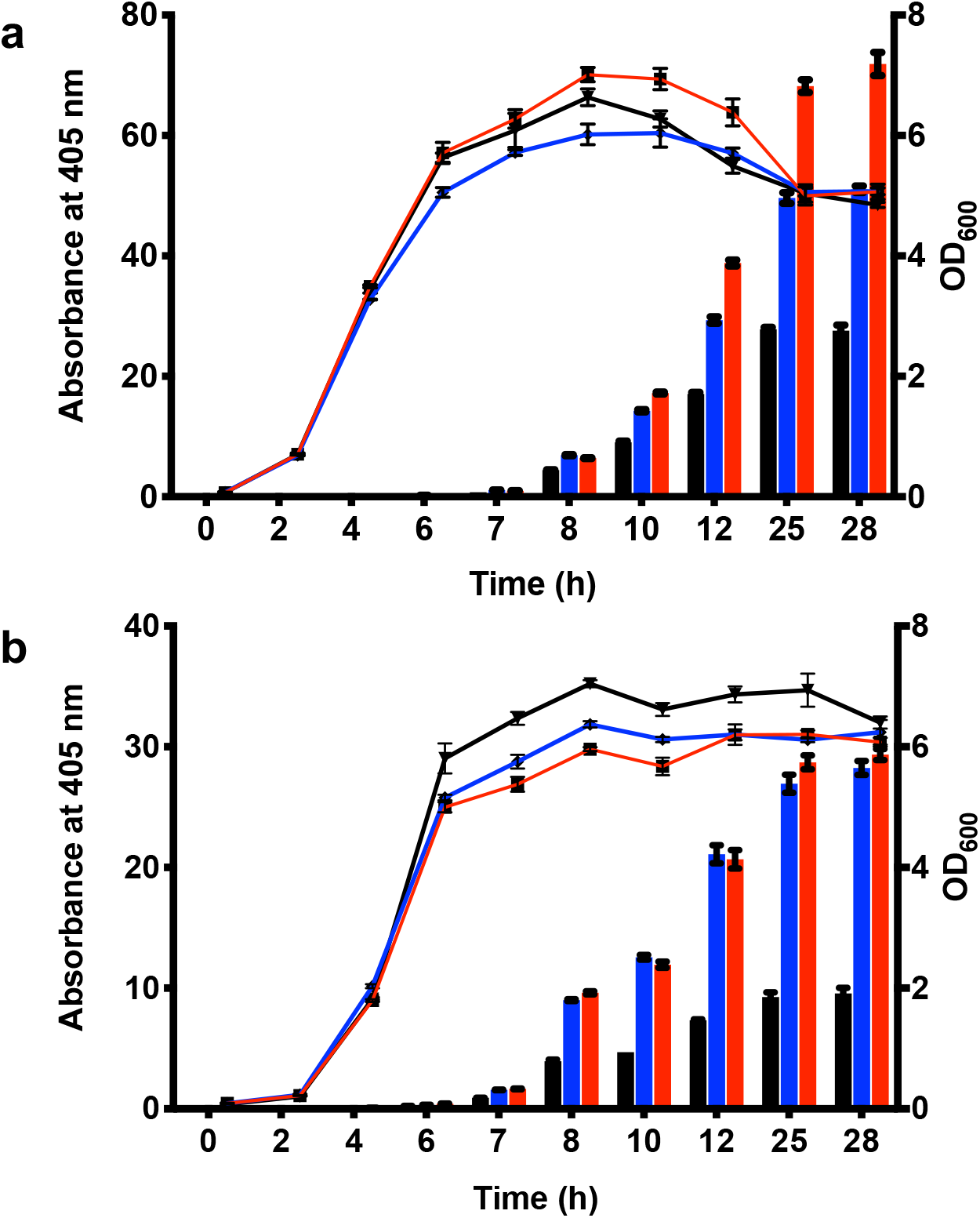
Protein secretion using OsmY(WT) and OsmY(M3) in *E. coli* C41 (DE3) and BL21(DE3). OsmY(WT)-Tfu0937 and OsmY(M3)-Tfu0937 were expressed using C41(DE3) and BL21(DE3) in 2×TY auto-induction media and at 37 °C and 250 rpm. Cultures were inoculated using 1:200 dilution. The cell optical densities monitored at 600 nm (right axis) are shown as lines. The activity of OsmY-Tfu0937 shown as bars was measured with the pNPG assay using an absorbance at 405 nm (left axis). The lines and bars of different variants are represented using different colours; OsmY(WT) – black, OsmY(TOA4) – blue and OsmY(M3) – red.

### OsmY(M3) variant improved the production of secretory RFP and Lipase 6B

Protein secretion is known to vary with the identity of the target protein. To determine if OsmY(M3) can be used to improve the production of other secretory proteins, the OsmY(WT) and OsmY(M3) were individually fused to the N-termini of the monomeric red fluorescent protein (RFP) and Lipase 6B. Secretion of RFP was determined by fluorescence intensity measurement of the spent medium (Fig. 8). Significant RFP secretion was observed after 24 h of cultivation (Fig. 8b), at which point OsmY(M3) showed more secretory RFP compared to the wildtype. The use of RFP allowed us to monitor intracellular protein easily (Fig. 8c). Comparing Fig. 8b and Fig. 8c helped shed light on the OsmY-assisted protein secretion. First, the protein production of OsmY(WT)-RFP was quicker compared to OsmY(M3)-RFP, as there was more total OsmY(WT)-RFP (intra- and extra-cellular) after 8 or 10 hours of cultivation. This was expected because the codons in OsmY(M3) were deoptimized. Second, extracellular protein was accumulated after 24 hours of cultivation (*i.e.*, stationary phase), consistent with Tfu0937 discussed above. This was true for both OsmY(WT) and OsmY(M3). The enhanced production of secretory RFP is contributed by two factors: (a) enhanced secretion [41% in OsmY (M3) vs 27% in OsmY(WT)] and (b) increased in total protein expression level (23% higher).

**Figure 8:**
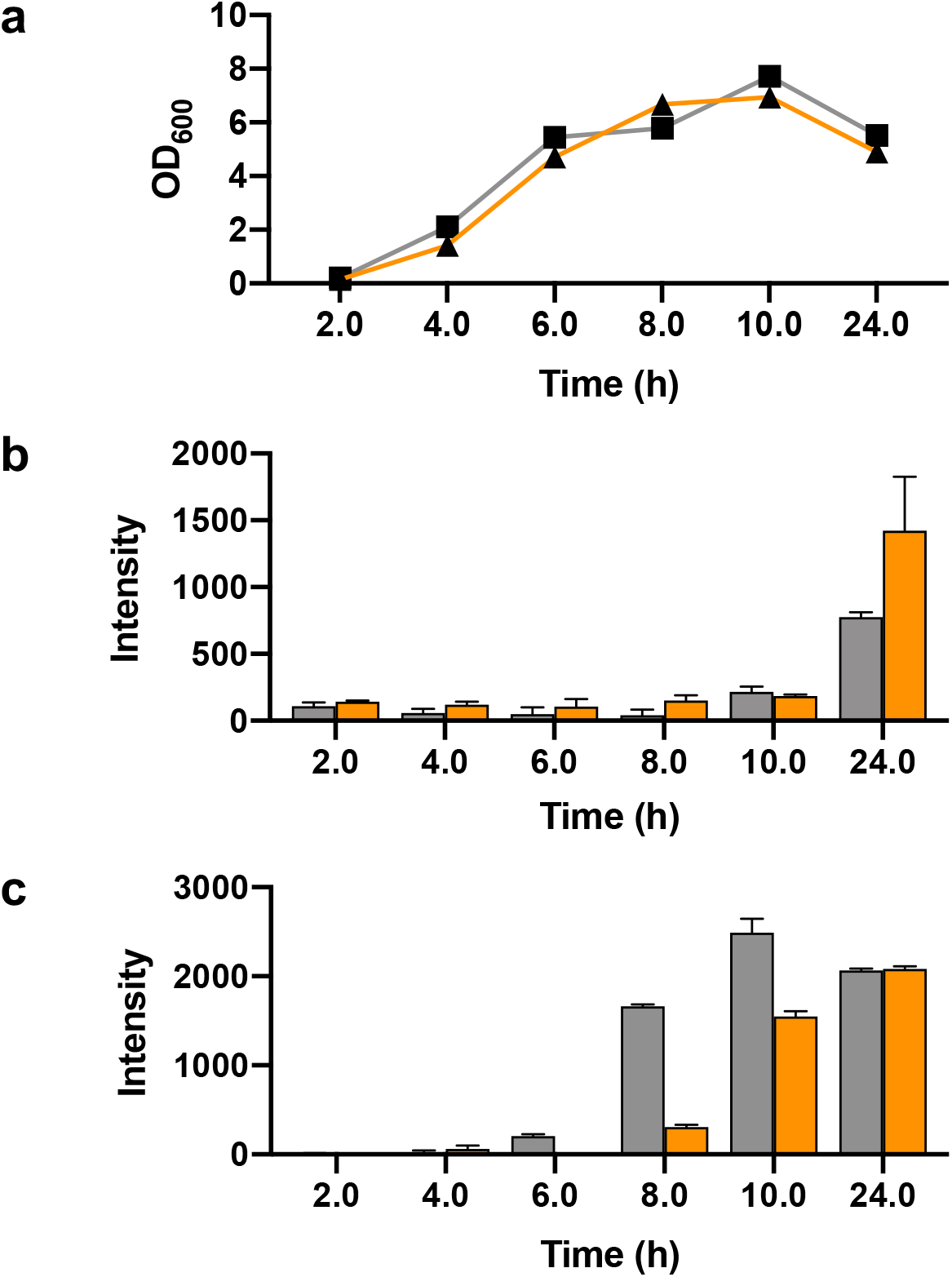
OsmY-RFP expression in *E. coli* BL21(DE3). Cells were cultivated in 20 mL of 2xTY auto-induction media at 37 °C using 1:100 dilution of inoculum. The supernatant and cells from 20 μL culture were used for fluorescence measurement using E_x_ = 584 nm and E_m_ = 607 nm. (**a**) Growth curve of the culture. (**b**) Fluorescence intensity of the spent medium. (**c**) Fluorescence intensity of the cells. The lines and bars of different variants are represented using different colours; OsmY(WT)-RFP – grey and OsmY(M3)-RFP – orange.

Lipase 6B is a thermostable variant of a lipase from *Bacillus subtillis* ^[28]^. OsmY(WT)-Lipase 6B and OsmY(M3)-Lipase 6B were expressed in both BL21(DE3) and C41(DE3), and the secreted protein was monitored using the *p*-nitrophenyl acetate (pNPA) assay (Fig. 9). The C41(DE3) strain grew faster than BL21(DE3) with increasing extracellular Lipase 6B activity detected after 7 h of cultivation. Variant OsmY(M3) produced 2.5 ± 0.3 times more secreted protein compared to OsmY(WT) after 10 h of cultivation, as determined by the pNPA assay. On the contrary, protein secretion was minimal in BL21(DE3) throughout the 10 h of cultivation monitored. Overexpression of recombinant lipase in *E. coli* can often lead to toxicity to the host ^[28, 29]^. The strain C41(DE3) was derived from BL21(DE3) and is commonly used for the expression of membrane proteins and toxic recombinant proteins ^[30]^, likely explaining its better performance for the production of secretory Lipase 6B.

**Figure 9:**
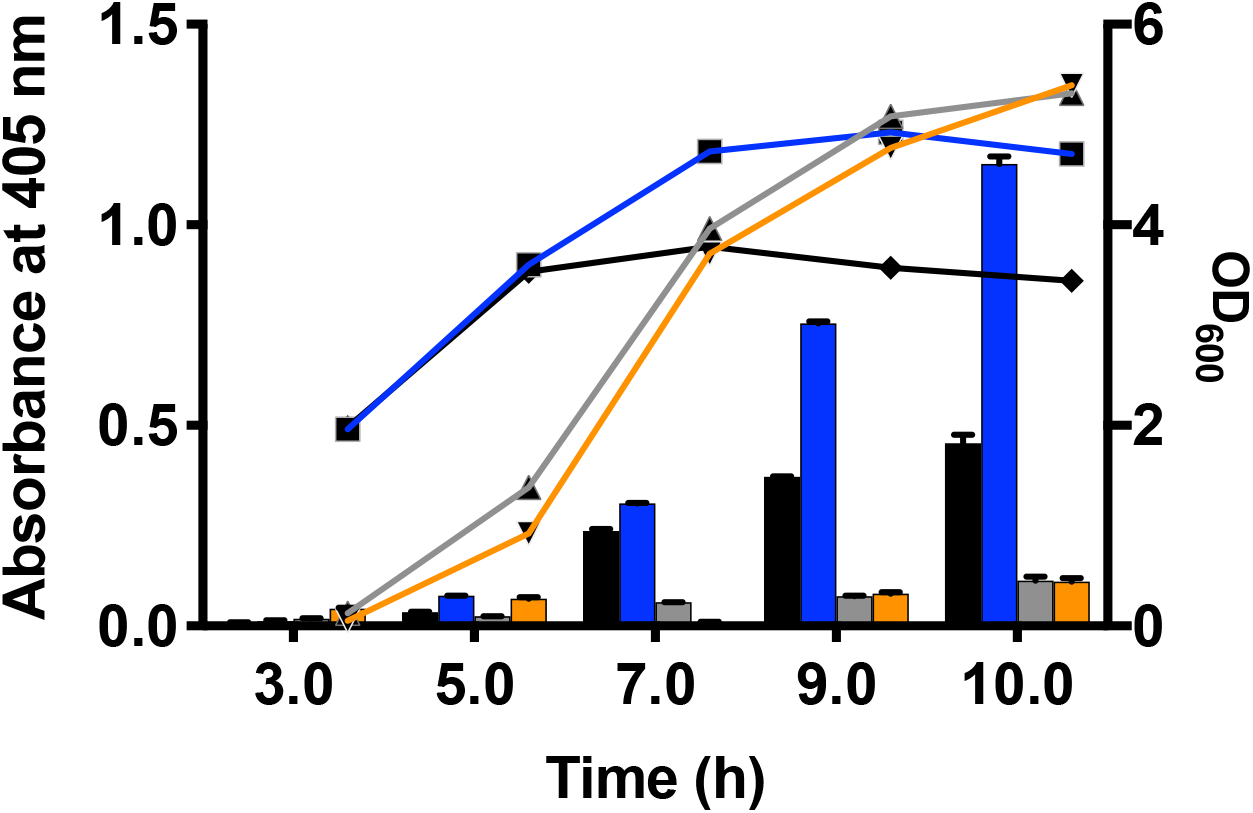
OsmY-Lipase 6B expressed in BL21(DE3) and C41(DE3). Cells were cultivated in 50 mL of 2xTY auto-induction media at 37 °C using 1:100 dilution of inoculum. The cell optical densities monitored at 600 nm are shown as lines. The activity of OsmY-Lipase6B in the spent medium measured with the pNPA assay using an absorbance at 405 nm are shown as bars. The lines and bars of different samples are represented using different colours; OsmY(WT)-Lipase 6B in C41(DE3) – black, OsmY(M3)-Lipase 6B in C41(DE3) – blue, OsmY(WT)-Lipase 6B in BL21(DE3) – grey and OsmY(M3)-Lipase 6B in BL21(DE3) – orange.

## CONCLUSIONS

In this study, we used directed evolution to improve the production of total secretory protein by OsmY. The best variant, OsmY(M3), enhanced the production of secreted β-glucosidase, Tfu0937, in two different *E. coli* strains by approximately 3 folds (Table 2). While directed evolution had been used to randomise the junction between the signal peptide and target protein ^[7]^ and to improve transporter proteins (*e.g.*, ABC transporter) ^[8]^, this is the first report that directed evolution of a secretory carrier was used to enhance secretory protein production in *E. coli*. The critical role of the signal peptide was reflected by the two important mutations (L6P ad S19C) found in the signal peptide. Importantly, this methodology can be further extended to evolve other signal peptide or carrier proteins for secretory protein production in *E. coli*.

**Table 2:** Production of secretory protein using OsmY(WT) and OsmY(M3) for three different target proteins.

| Target protein | M<sub>w</sub> (kDa) | Theoretical pI | Protein source organism | Total secretory protein production by OsmY(M3) relative to OsmY(WT) / cultivation time (h) | Recombinant protein expression host |
| --- | --- | --- | --- | --- | --- |
| Tfu0937 | 53.4 | 4.81 | *Thermobifida fusca* YX | 3.1 ± 0.3 / 25 | BL21(DE3) |
|  |  |  |  | 2.9 ± 0.8 / 24 | C41(DE3) |
| RFP | 25.4 | 5.65 | Synthetic construct | 1.8 ± 0.6 / 24 | BL21(DE3) |
| Lipase 6B | 19.7 | 6.96 | *Bacillus subtilis* 168 | 2.5 ± 0.3 / 10 | C41(DE3) |

Subcellular fractionation of Tfu0937 with and without OsmY fusion showed that OsmY was essential for the extracellular secretion and that >80% of OsmY-Tfu0937 protein was found extracellularly after 15 h of cultivation (Fig. 4). The transferability of improved secretory protein production by OsmY(M3) to a different *E. coli* stain and for other proteins, RFP and Lipase 6B (Table 2), showed the versatility of the OsmY(M3) variant identified. It is worth noting that these are three proteins originating from diverse organisms, having diverse molecular weights (19.7 – 53.4 kDa) and protein functions (Table 2). Together with previous studies on OsmY as a secretory carrier ^[17–21]^, this work demonstrated the potential application of OsmY and its variants for secretion of a broad range of proteins.

## METHODS

### Materials

Chemicals were purchased from Sigma-Aldrich (Dorset, UK) and ForMedium (Norfolk, UK). DNA modifying enzymes, deoxyribonucleotides, and DNA ladders were purchased from New England Biolabs (Hitchin, UK), Thermo Fisher Scientific (Loughborough, UK), and Agilent Technologies (Cheadle, UK). Nucleic acid purification kits were purchased from Machery-Nagel (Düren, Germany) and Omega Bio-tek (Norcross, USA). All oligonucleotides were synthesized by Eurofins Genomics (Ebersberg, Germany), and summarized in Table S1.

### Strains

*Escherichia coli* DH5α was used for all molecular cloning, plasmid propagation and maintenance. *E. coli* BL21 (DE3) (Merck; Darmstadt, Germany) and C41(DE3) (Lucigen; Wisconsin, USA) were used for protein expression.

### Molecular cloning of Tfu0937, OsmY-Tfu0937, OsmY-RFP and OsmY-Lipase 6B

The DNA sequences encoding the *E. coli* osmotically-inducible protein Y (OsmY; GenBank: AUY30809.1), the *Thermobifida fusca* YX β-glucosidase (Tfu0937; GenBank: AAZ54975.1), the *Bacillus subtilis* lipase mutant (Lipase 6B; ^[28]^), and a monomeric red fluorescent protein (RFP, GenBank: AAM54544.1) were codon-optimized for protein expression in *E. coli* and synthesized by GenScript (Piscataway, USA). The OsmY-Tfu0937 gene (supplementary Fig. S1b) was cloned into pET-24a(+) vector (Merck; Darmstadt, Germany) using NdeI and EcoRI sites, and the resulting plasmid (pET24a-OsmY-Tfu0937; Fig. S1a) was used for protein engineering in this study.

To create the pET24a-Tfu0937 plasmid, the Tfu0937 gene was amplified with NdeI-Tfu0937-F and EcoRI-Tfu0937-R primers, digested with NdeI and EcoRI, and cloned into the pET-24a(+) vector using NdeI and EcoRI sites. To create pET24a-OsmY-RFP and pET24a-OsmY-Lipase 6B, the RFP gene was amplified using primers BamHI-RFP-F and EcoRI-RFP-R and the Lipase 6B gene was amplified using primers BamHI-Lipase6B-F and Lipase6B-EcoRI-R. The amplified RFP and Lipase 6B genes were digested with BamHI and EcoRI, and cloned into the BamHI/EcoRI digested pET24a-OsmY-Tfu0937 plasmid to replace the Tfu0937 gene. The sequences of all primers were summarised in Table S1.

### Random mutagenesis by epPCR

Two error-prone polymerase chain reaction (epPCR) conditions were used in this study: high and low mutagenic rates. For epPCR of high mutagenic rate, the 50-μL PCR mixture contained 1× standard Taq reaction buffer (Mg-free), 7 mM MgCl_2_, 0.05 mM MnCl_2_, 0.2 mM dATP, 0.2 mM dGTP, 1 mM dTTP, 1 mM dCTP, 20 pmol OsmY-DirEv-F primer, 20 pmol OsmY-DirEv-R primer (Table S1), 50 ng pET-24a(+)-OsmY-Tfu0937 and 1.25 U Taq DNA polymerase (New England Biolabs). For epPCR of low error rate, the 50-μL PCR mixture contained 1×standard Taq reaction buffer (Mg-free), 1.5 mM MgCl_2_, 0.01 mM MnCl_2_, 0.3 mM of each dNTP, 4.5 pmol OsmY-DirEv-F primer, 4.5 pmol OsmY-DirEv-R (Table S1), 7.77 ng pET-24a(+)-OsmY-Tfu0937 and 1.25 U Taq DNA polymerase (New England Biolabs). The PCR mixtures were thermocycled using the following conditions: (i) 30 s initial denaturation at 95 °C, (ii) 25 cycles of 20 s denaturation at 95 °C, 30 s annealing at 52 °C and 1 min 30 s extension at 68 °C, and (iii) 5 min final extension at 68 °C. PCR products were purified by gel extraction. After restrictive digestion with NdeI and BamHI, the PCR products were cloned into NdeI/BamHI digested pET24a-OsmY-Tfu0937 plasmid. The recombinant plasmids were subsequently electroporated into *E. coli* C41(DE3) and plated on TYE agar (10 g/L tryptone, 5 g/L yeast extract, 8 g/L NaCl and 15 g/L agar) supplemented with 50 μg/mL kanamycin to create the OsmY mutant libraries.

### Site directed mutagenesis to create variants OsmY(L6P), OsmY(S154R) and OsmY(V191E)

Site directed mutagenesis experiments were performed using partially overlapping primers designed by the OneClick program ^[31]^. Primers OsmY(L6P)-FP and OsmY(L6P)-RP were used for OsmY(L6P), primers OsmY(V154R)-FP and OsmY(S154R)-RP were used for OsmY(S154R), and primers OsmY(V191E)-FP and OsmY(V191E)-RP were used for OsmY(V191E) (Table S1). In the two-stage PCR designed by OneClick ^[31]^, PfuUltra High-Fidelity polymerase (Agilent Technologies) was used to introduce the L6P and S154R mutations, and PfuUltra II Fusion HS DNA Polymerase (Agilent Technologies) was used to introduce the V191E mutation.

### Cultivation and protein expression in 96-well microplate

Individual colonies of the mutant library were manually transferred from the agar plates using sterile toothpicks into 96-well deep-well microplates, with each well containing 1.5 mL 2×TY medium (16 g/L tryptone, 10 g/L yeast extract and 5 g/L NaCl) supplemented with 50 μg/mL kanamycin. Column 6 on the microplate was inoculated with the wildtype as internal control. These master plates were covered with lids, sealed and cultivated at 30 °C for 24 h.

To express protein for screening, master plates were replicated using a pin replicator into fresh 96-well deep-well plates, with each well containing 1.5 mL 2×TY auto induction medium [16 g/L tryptone, 10 g/L yeast extract, 3.3 g/L (NH_4_)_2_SO_4_, 6.8 g/L KH_2_PO_4_, 7.1 g/L Na_2_HPO_4_, 0.5 g/L glucose, 2.0 g/L α-lactose and 0.15 g/L MgSO_4_] supplemented with 50 μg/mL kanamycin. The plates were cultivated at 30 °C for 24 h before (a) they were transferred to a fridge and kept at 4 °C overnight for the cells to settle or (b) they were plate centrifuged at 4000 rpm for 10 min (Heraeus Megafuge 8R; M10 microplate rotor). The supernatant containing the secreted OsmY-Tfu0937 was used for high-throughput screening on the next day.

Abgene 96-well polypropylene deep-well plates (Thermo Fisher Scientific; AB0661) and Abgene polypropylene plate covers (Thermo Fisher Scientific; AB0755) were used in preparing master plates and protein expression plates. All plate cultivations were conducted in Titramax 1000 plate shaker coupled to an Incubator 1000 heating module (Heidolph Instruments; Essex, UK) using a shaking speed of 1050 rpm.

### High-throughput screening in 96-well microplate

Flat-bottom clear 96-well polystyrene microplates (Greiner Bio-One; 655161) were used for screening. Unless otherwise stated, 20 μL of spent medium was transferred to 96-well microplates, with each well containing 30 μL of 0.2 M sodium acetate. Fifty μL of pNPG solution (5.3 mM pNPG, 0.2 M sodium acetate, pH 4.8) was then added to each well. The microplates were incubated at 50 °C and 1050 rpm in a microplate shaker for 5 min before 100 μL of NaOH-Gly buffer (0.4 M glycine, pH 10.8) was added to stop the reaction. Absorbance at 405 nm was recorded with Multiskan FC microplate photometer (Thermo Fisher Scientific). All shaking steps were conducted in Titramax 1000 (Heidolph Instruments) using a shaking speed of 1050 rpm.

### Variant characterization using tube and flask cultures

The variants were characterized using two methods, (a) tube cultures and (b) flask cultures. For tube cultures, 5 μL of cells from the master plate was transferred to culture tubes containing 5 mL of 2xTY auto induction medium supplemented with 50 μg/mL kanamycin and cultivated overnight at 37 °C. Flask cultures were used for expression studies, where 20 mL or 50 mL of 2×TY-based auto induction medium was inoculated using 1:100 or 1:200 dilution of the pre-culture. Cells were then cultivated at 37 °C and 240 rpm, unless otherwise stated. During cultivation, cell growth was monitored by absorbance reading at 600 nm (OD_600_). For assay measurement, 0.5 mL to 1.0 mL of culture was sampled and secreted protein was measured using the pNPG assay (Tfu0937), the pNPA assay (Lipase 6B) or fluorescence intensity (RFP).

### The pNPG assay, pNPA assay and RFP fluorescence measurement

The sampled culture (0.5–1.0 mL) was centrifuged at >17000 g for 5–10 min and 50 μL of the supernatant containing the secreted protein was used for assay. The supernatant was diluted to keep within the assay linear detection range when necessary. For pNPG assay, 50 μL of supernatant was transferred to a microplate and sample was assayed using the same protocol as high-throughput screening. The supernatant was diluted using either 2×TY medium or sample buffers for the subcellular fractions when necessary. For pNPA assay, 20 μL of supernatant was transferred to a microplate and 180 μL of freshly prepared pNPA solution (2.0 mM pNPA, 50 mM K_x_PO_4_, pH 7.2) was added. Reaction was incubated at room temperature for 10 min before the absorbance was recorded at 405 nm. The supernatant was diluted using 50 mM K_x_PO_4_, pH 7.2 assay buffer when necessary. For RFP detection, 100 μL of supernatant was transfer to a black microplate (Greiner Bio-One; 655906) and fluorescence intensity was measured using the SpectraMax M2e plate reader (Molecular Device; California, USA) at excitation wavelength of 584 nm and emission wavelength of 607 nm. The RFP fluorescence intensity of the cell was measured using cells from 100 μL of culture resuspended in 100 μL of 50 mM K_x_PO_4_, pH 7.5 and at the same excitation and emission wavelengths. Assay background, for the pNPG and pNPA assays, was always removed by subtracting the assayed absorbance (at 405 nm) of the spent medium from the culture of cells without plasmids [*i.e.*, BL21(DE3) or C41(DE3)].

### Soluble protein extraction using enzymatic lysis

One mL of culture was centrifuged at >17000 g for 5–10 min. The supernatant was removed, and the cell pellet was resuspended in 200 μL of lysomix (10 mg/mL lysozyme in 50 mM, pH 7.5) by vortexing. Sample was incubated on ice for 1 h followed by centrifugation at >17000 g for 10 min. The supernatant, which contained the soluble protein fraction, was transferred to a fresh tube. OsmY-Tfu0937 activity in the soluble fraction was determined using the pNPG assay.

### Subcellular fractionation of Tfu0937 and OsmY-Tfu0937

Cells were cultivated as described earlier using 50 mL or 100 mL of 2×TY-based auto induction medium. The cells were harvested by centrifugation at 6000 g, 4 °C for 5 min. The supernatant was transfer to a clean tube and used as the extracellular fraction. The cell pellet was resuspended in 25 mL of osmotic shock buffer (100 mM Tris–HCl, 20% sucrose, 1 mM EDTA, pH 8.0) and incubated on ice for 15 min. The cell suspension was then centrifuged at 7000 g, 4 °C for 10-15 min and the clear supernatant was transferred to a clean tube (fraction P1). The resulting cell pellet was resuspended in 2.5 mL of ice-cold water with 1 mM MgCl_2_ and incubated on ice for 15 min. The cell suspension was centrifuged at 7000 g, 4 °C for 10 min and the clear supernatant was transferred to a clean tube (fraction P2). Fractions P1 and P2 were used to determine Tfu0937 activity in the periplasmic fraction. The cell pellet following periplasmic extraction was resuspended in 10 mL of 100 mM Tris-HCl buffer (pH 8.0) by vortexing and cells were lysed by sonication (Vibra-Cell ultrasonic liquid processors; Sonics & Materials; Newtown, USA) using 40% amplitude and 1.5 min total processing time. After sonication, 2 mL of sonicated sample was centrifuged at 20000 g, 4 °C for 10 min. The supernatant was collected as the cytoplasmic fraction.

## LIST OF ABBREVIATIONS

dATP: deoxyadenosine triphosphate
dCTP: deoxycytidine triphosphate
dGTP: deoxyguanosine triphosphate
dNTP: deoxynucleotide
dTTP: deoxythymidine triphosphate
DyP4: dye-decolorizing peroxidase 4 from *Pleurotus ostreatus* strain PC15
E_m_: emission wavelength
*E. coli*: *Escherichia coli*
E_x_: excitation wavelength
epPCR: error-prone polymerase chain reaction
HTS: high-throughput screening
M_w_: molecular weight
pNPA: *p*-nitrophenyl acetate
pNPG: *p*-nitrophenyl-β-D-glucopyranoside
OD_600_: optical density at 600 nm
OsmY: osmotically-inducible protein Y
PCR: polymerase chain reaction
RFP: red fluorescent protein
WT: wildtype

## ADDITIONAL INFORMATION

## Acknowledgement

We thank EPSRC (EP/E036252/1), the Open Project Funding of the State Key Laboratory of Bioreactor Engineering (to JX and TSW), COST action (CM1303 Systems Biocatalysis; to DGP), the Royal Academy of Engineering (the Leverhulme Trust Senior Research Fellowship; to TSW; LTSRF1819\15\21), BBSRC GCRF-STARS (BB/R020183/1; to TSW), The University of Sheffield (GCRF Fellowship; to KLT), the Department of Chemical and Biological Engineering (Summer Undergraduate Fellowship; to JR, MCMW and YPY) for financial support.

## Authors’ contributions

KLT and TSW supervised the project and designed the experiments. KLT, JR, DGP, SKT, MCMW, YPY and NN conducted the experiments. KLT, TSW and JX analysed the data and wrote the manuscript.

## Competing interests

The author(s) declare no competing interests.

## Availability of data and material

All data generated or analysed during this study are included in this published article and its supplementary information files.

## Ethics approval and consent to participate

Not applicable.

## Consent for publication

All authors have read the manuscript and approved its submission.

## Supplementary material

**Table S1:**
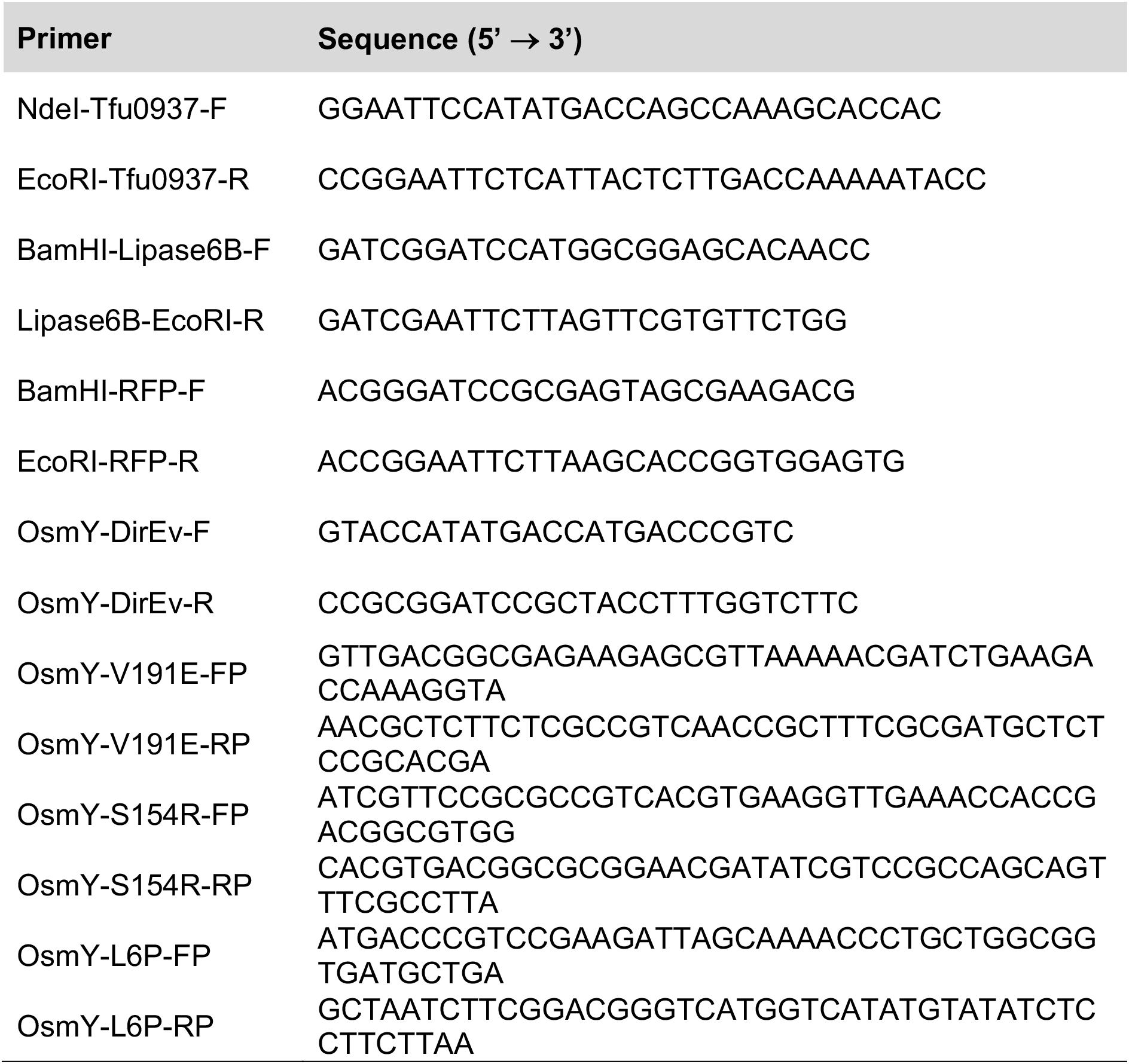
Primers used in this study

**Figure S1:**
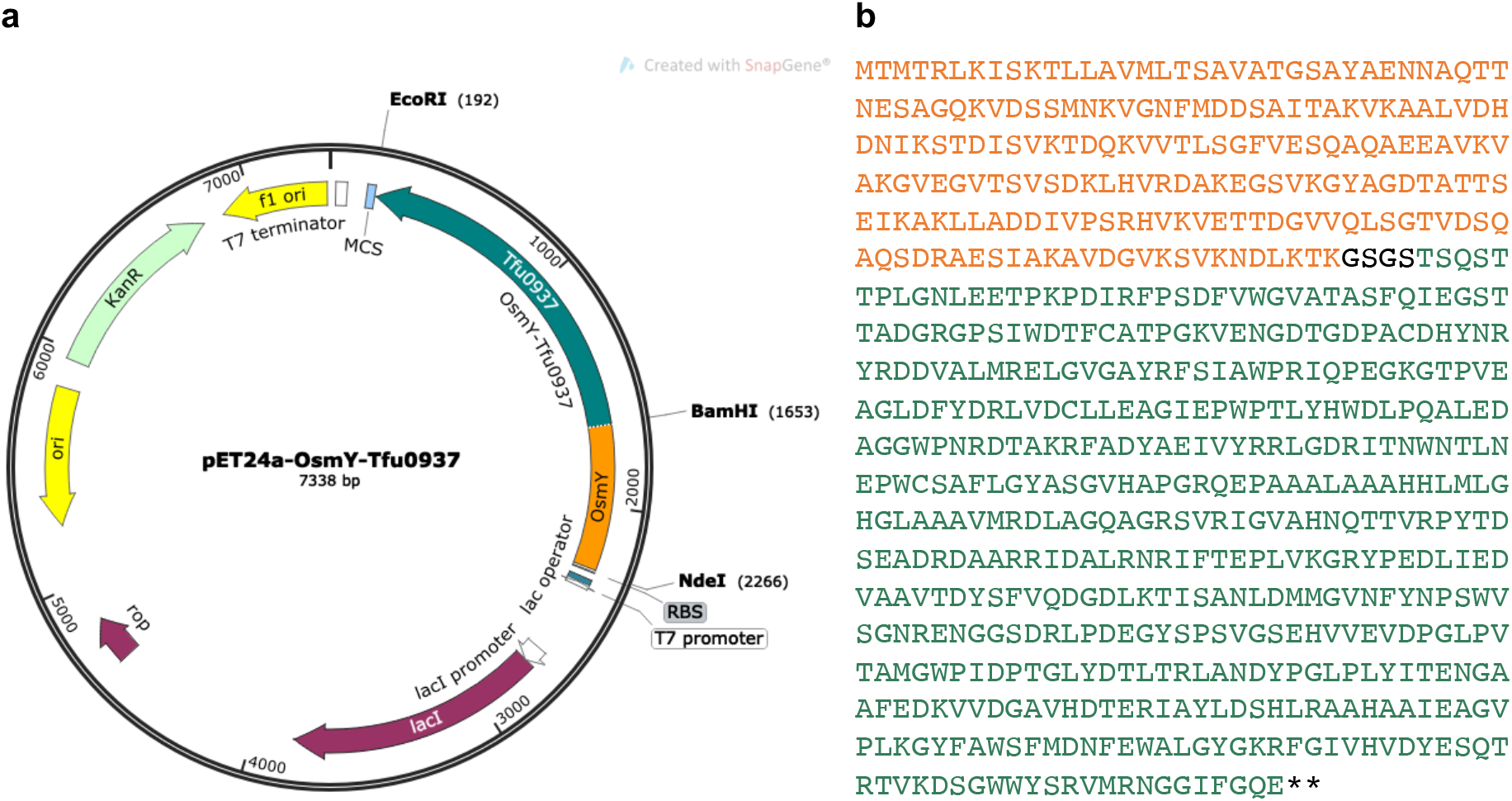
**(a)** Plasmid map of pET24a-OsmY-Tfu0937. **(b)** Amino acid sequence of OsmY-Tfu0937. The OsmY wildtype sequence is shown in orange, Tfu0937 sequence in green, and GSGS linker sequence in black.

**Figure S2:**
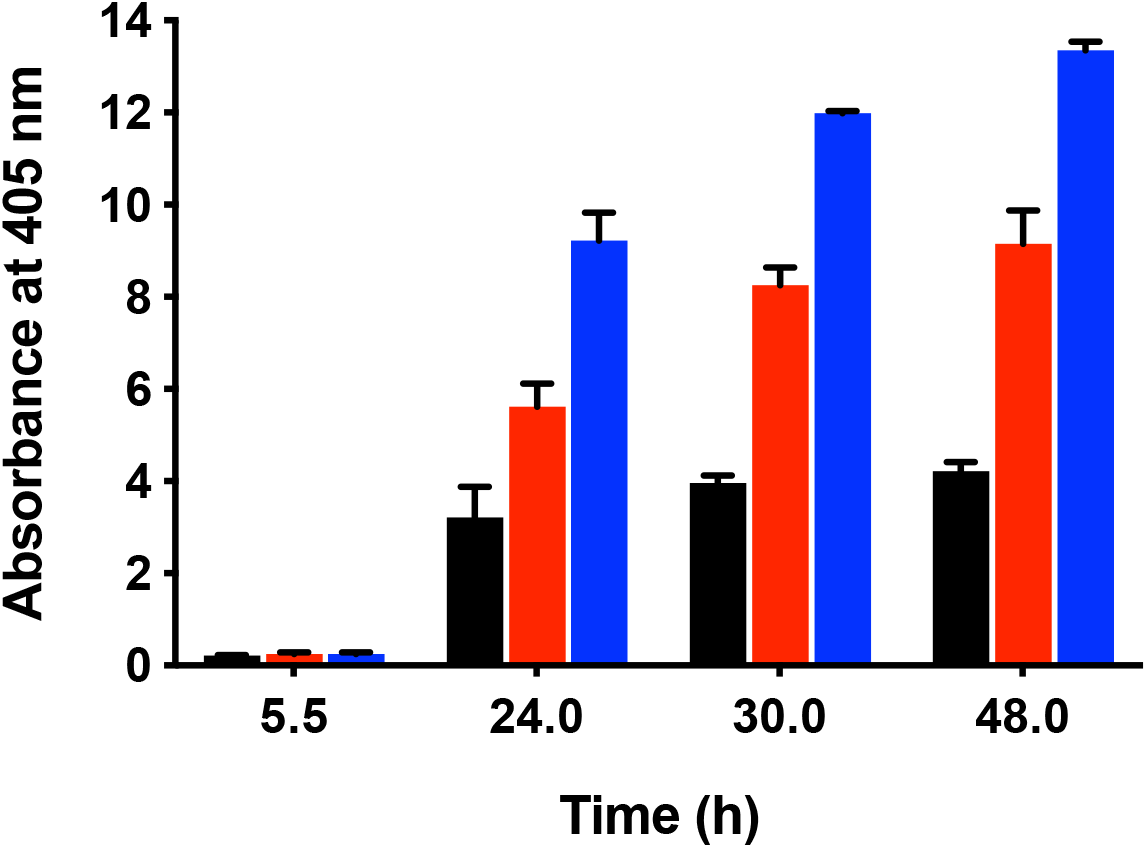
Characterization of improved variants in 50-mL flask cultures. The activity of extracellular OsmY-Tfu0937 was determined with the pNPG assay using an absorbance at 405 nm. Cultures were inoculated using 1:100 dilution. OsmY(WT) type and variants were expressed using C41(DE3) in 2TY auto-induction media and at 37 °C and 200 rpm. The activities of wildtype and variants are represented using different colours; OsmY(WT) (black), OsmY(TOA4) (red) and OsmY(M3) (blue). Experiments were performed in triplicates.

**Figure S3:**
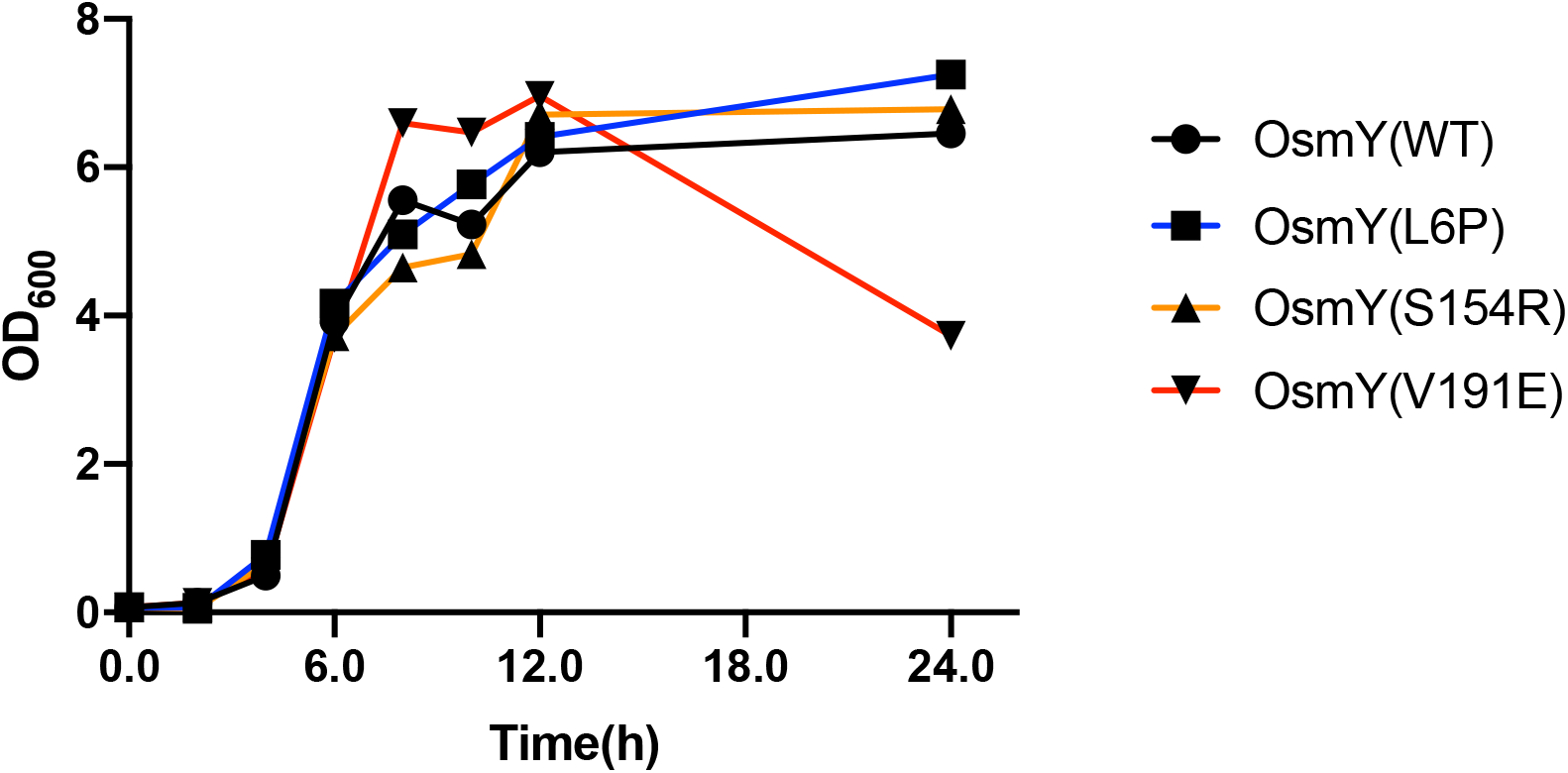
Growth curve of single site-directed mutants measured at 600 nm. Cells were cultivated in 20-mL flask cultures using 2×TY auto-induction media at 37 °C. Culture was inoculated using 1:100 dilution and 0.5 mL culture was sampled at various time points for cell density (OD_600_) measurement. Wildtype and variants are represented using different colours; OsmY(WT) – black line, OsmY(L6P) – blue line, OsmY(S154R) – orange line and OsmY(V191E) – red line.

## Notes

### Competing Interest Statement

The authors have declared no competing interest.

